# Competition and Caries on Enamel of a Dual-species Biofilm Model of *Streptococcus mutans* and *Streptococcus sanguinis*

**DOI:** 10.1101/2020.05.28.122697

**Authors:** Natalia Díaz-Garrido, Carla P. Lozano, Jens Kreth, Rodrigo A. Giacaman

## Abstract

Imbalances within the dental biofilm trigger dental caries, currently considered a dysbiosis and the most prevalent non-communicable disease. There is still a gap in knowledge about the dynamics of enamel colonization by bacteria from the dental biofilm in caries. The aim, therefore, was to test whether the sequence of enamel colonization by a typically commensal and a cariogenic species modifies biofilm’s cariogenicity. Dual-species biofilms of *Streptococcus mutans* (Sm) and *Streptococcus sanguinis* (Ss) on saliva-coated enamel slabs were inoculated in different sequences: Sm followed by Ss (Sm-Ss), Ss followed by Sm (Ss-Sm), Sm and Ss inoculated at the same time (Sm=Ss) and the single-species controls Sm followed by Sm (Sm-Sm) and Ss followed by Ss (Ss-Ss). Biofilms were exposed to 10% sucrose, 3x/day for 5 days and the slabs/biofilms were retrieved to assess demineralization, viable cells, biomass, proteins, polysaccharides and H_2_O_2_ production. When compared with Sm-Sm, primary inoculation with Ss reduced demineralization (p<0.05). Both Ss-Sm and Sm=Ss sequences showed reduction in biomass, protein and polysaccharide content (p<0.05). The highest *S. sanguinis* viable cells and H_2_O_2_ production and the lowest acidogenicity were observed when Ss colonized enamel before Sm (p<0.05). Initial enamel adherence with commensal biofilms seems to induce more intense competition against more typically cariogenic species, reducing cariogenicity.

**Importance:** The concept of caries as an ecological disease implies the understanding of the intricate relationships among the populating microorganisms. Under frequent sugars exposure, some the bacteria from the oral biofilm develop pathogenic traits that lead to oral imbalances, known as dysbiosis. Depending on which microorganism colonizes the dental surface first, different competition strategies may be developed. Since the study of the interactions in the entire dental biofilm is not an easy task, in this article we model the interplay among these microorganisms using a caries-inducing (*S. mutans*) and a health-associated species (*S. sanguinis*). Initial enamel adherence with S. sanguinis seems to induce more intense competition against more typically caries-inducing species. Besides continuous exposure with sugars, early colonization of the enamel by highly cariogenic species, like *S. mutans*, appears to be needed to develop caries lesions, as well. Promoting early colonization by health-associated bacteria, such as *S. sanguinis*, could help maintaining oral health, delaying dysbiosis.

## Introduction

Dental caries and periodontal diseases have been defined as microbial dysbiosis (1), but the role played by each constituent of the multispecies microbial biofilm is far from being fully understood. It has been recognized that commensal streptococci act as early colonizers of the enamel (2), binding other early colonizers and host molecules to initiate the dental biofilm formation. *Streptococcus sanguinis* (*S. sanguinis*) is a commensal member of the early colonizers in the dental biofilm that has been more abundantly recovered in caries-free children (3) and adults (4). Conversely, another important oral streptococcus, *Streptococcus mutans* (*S. mutans*), is not considered an early colonizer, but is endowed with a powerful machinery to metabolize carbohydrates, producing critical amounts of acids as well as efficiently generating an adherent extracellular polysaccharide matrix implicated in caries development (5). An inverse relation between *S. mutans* and *S. sanguinis* counts has been described (6), so when high number of colonies of *S. mutans* are recovered from the biofilm, relatively lower numbers of *S. sanguinis* are obtained. This opposite trend suggest competition between both species. Drivers of competition between both species are nutrient availability or fitness within the ecological niche (7–9). Each species has developed strategies to mutually inhibit each other (10). Hence, *S. mutans* can produce bacteriocins (mutacins), which are used to inhibit competing species, including *S. sanguinis* (9, 11). On the other hand, *S. sanguinis* produces hydrogen peroxide (H_2_O_2_; encoded by the *spxB* gene (12), as an antimicrobial compound, which during the early stages of biofilm formation is a powerful tool to exclude competing species, as peroxides are toxic for bacteria like *S. mutans* (13).

Dental caries is a disease characterized by lactic acid-induced hard dental tissue demineralization, caused by frequent carbohydrate exposure to the dental biofilm, which shifts the ecological balance towards a non-infectious polymicrobial dysbiosis (14). Despite the existence of evidence from clinical studies on the interacting relationship between commensal and cariogenic bacteria within the dental biofilm, the effect of the order in which they adhere to the enamel under environmental stressors relevant for the caries process, such as frequent sucrose exposure, has not been reported. Understanding whether primary colonization of the dental tissues by cariogenic or by commensal microorganisms, promotes competition between them and whether this competition modifies the structure and functionality of biofilm on the hard-dental tissue, is of interest and has not been characterized in a caries model with dual-species biofilms. The aim of the study was, therefore, to test if the sequence of enamel adherence (colonization) by *S. sanguinis* and *S. mutans* modifies resulting cariogenicity.

## Materials and Methods

### Enamel slab preparation and acquired pellicle formation

Based on an established single-species caries model with biofilms of *S. mutans* (15), a dual-species caries model was applied. Dental enamel slabs were prepared from bovine incisors, as described (15) and autoclaved. Slabs were mounted on metal brackets made with orthodontic wire and suspended into in the wells of a 24-well plate (Costar®, Corning, NY, USA). Slabs were covered with ultrafiltered saliva from two healthy donors for 30 min to stimulate the formation of an acquired pellicle-like layer.

### Formation of single and dual-species biofilms of *S. mutans* and *S. sanguinis*

Frozen stocks of *S. mutans* UA159 (isolated from a child with active caries and kindly donated by Prof. J.A. Cury, UNICAMP, Brazil) and *S. sanguinis* SK36 (originally isolated from human dental plaque and donated by J. Kreth) were reactivated in brain heart infusion broth (BHI; Merck, Darmstadt, Germany) supplemented with 1% glucose and incubated at 37°C and 10% CO_2_ (Panasonic, MCO-19M, Osaka, Japan) for 18 h. The optical density (OD_600_) was adjusted to 0.1 (corresponding to 10^3-4^ CFU/mL). A culture aliquot of 100 μL from each species was inoculated onto acquired pellicle-covered slabs with BHI medium supplemented with 1% sucrose to form adherent biofilms (16). To characterize the results of sequential colonization of enamel, the following inoculation sequences were assayed; (1) *S. mutans* followed by *S. mutans* (Sm-Sm) (control), (2) *S. sanguinis* followed by *S. sanguinis* (Ss-Ss) (control), (3) *S. mutans* followed by *S. sanguinis* (Sm-Ss), (4) *S. sanguinis* followed by *S. mutans* (Ss-Sm) and (5) both species at the same time (Ss=Sm). Due to differences in biofilm formation, Ss biofilms were allowed to grow for 16 h, before Sm was inoculated, whereas Sm biofilms were allowed to grow for 8 h before Ss was inoculated. Subsequently, to mimic salivary basal glucose concentration, biofilms were allowed to mature in BHI medium supplemented with 0.1 mM glucose, for 24 h (17).

### Sucrose exposure

For 5 days, slabs/biofilms were exposed 3 times per day to 10% sucrose for 5 min, washed 3 times with 0.9% NaCl and returned to a plate with BHI supplemented with 0.1 mM glucose. Culture medium was replaced twice per day, before the first and after the last exposure to sucrose. The caries-negative control was, instead, exposed to 0.9% NaCl for 5 min, with the same regime. Two independent experiments in triplicate were carried out (n=6). The initial phase to promote adhesion and biofilm formation was carried out with BHI + 1% sucrose, but before and during the cyclic exposures to sucrose, enamel slabs/biofilms were grown only in BHI with 0.1 mM glucose. Enamel slabs/biofilms were never simultaneously exposed to glucose and sucrose.

### Biofilm acidogenicity

To monitor acid production, medium pH was measured with a microelectrode (Orion 910500, Thermo Scientific, Waltham, MA, USA) coupled to a pH-meter (Orion Star A211, Thermo Scientific). Individual measurements were made twice per day, after each medium change.

### Enamel demineralization assessment

The percentage of surface Knoop microhardness loss (%SHL) was performed (18). Before the experiments, the initial surface microhardness (SH_i_) of the enamel slabs was determined. After completion of the 5 days experimental period, slabs were mounted on a glass plate, and a second SH measurement was obtained, considered as final (SHf) (kg/mm^-2^). Each SH test was performed with three indentations separated by 100 μm each. Mean values for SH_i_ and SH_f_ were used to calculate the %SHL: (SH_i_ average - SH_f_ average) x 100/ SH_i_ average.

### Biofilm analysis

After completion of the experiments, slabs were washed and homogenized in 0.9% NaCl for 30s (Maxi Mix II type 37600 Mixer, Thermolyne, Iowa, USA), which causes biofilm detachment (18). Biofilm suspensions were saved to evaluate biomass, viable microorganisms, insoluble extracellular polysaccharide formation, total protein content and H_2_O_2_ production, all based on previously described methods, so just a brief description follows below.

### Biomass

The dry weight of the samples was used to determine the biomass (16). A volume of 200 μL of the biofilm suspension was transferred to a previously weighed tube (W_i_) and incubated with absolute ethanol at −20°C for 15 min. The pellet was dried by liquid evaporation at 37°C for 24 h to obtain the final dry weight (W_f_). To obtain the biomass, the following formula was applied: W_i_-W_f_, normalized to mg/mL of biofilm suspension.

### Protein content of the biofilm

A 50 μL aliquot of the biofilm suspension was treated with 2M NaOH and incubated at 100°C for 15 min (17). The supernatant was used to determine the total protein concentration by the Bradford method (Bradford reagent, Merck, Darmstadt, Germany), in a microplate reader at 595 nm. Results were expressed as μg/mg of biomass.

### Insoluble extracellular polysaccharide (IEPS) formation

(19) A 200 μL aliquot of the biofilm suspension was centrifuged and the resulting pellet was treated with 200 μL of 1M NaOH, homogenized and centrifuged again. The pellet was treated with three volumes of cold absolute ethanol and the pellet was washed with 70% cold ethanol and centrifuged again. The pellet was resuspended in 1M NaOH and total carbohydrates concentration was obtained by the sulfuric phenol method (20). Results were normalized by dry weight and expressed as percentage of polysaccharides by mg of biomass.

### Counts of viable cells

A 100 μL aliquot of the biofilm suspension was serially diluted up to 1: 10^8^ (v/v) in 0.9% NaCl. A drop of 50 μL of each dilution was seeded on Prussian blue agar (21) for *S. sanguinis* and Mitis Salivarius agar (Difco, BD, New Jersey, USA) supplemented with 0.2 units/mL of bacitracin for *S. mutans*, both in triplicate. After incubation for 48 h, phenotypic observation and counting was carried out for each plate under magnification (4x) and the number of colonies, corrected by the dilution factor were normalized by biomass dry weight and expressed as CFU/mL.

### H_2_O_2_ production

To assess peroxide production (22), the supernatants from the single-species and dual-species biofilm cultures, at the end of the experiments, were recovered and centrifuged. Resulting pellets were resuspended in 1 mL of BHI, centrifuged and the supernatant was filtered. The amount of H_2_O_2_ was obtained using the Amplex®, Red Hydrogen Peroxide/Peroxidase Assay kit (Molecular Probes, Invitrogen, Burlington, Ontario, Canada).

### Statistical analysis

Data were analyzed using the statistical software SPSS v15.0 for Windows (SPSS Inc, Chicago, USA). The variables acidogenicity, demineralization, biomass, total proteins, insoluble extracellular polysaccharides, viable microorganisms and H_2_O_2_ production were analyzed using a multiple comparison by ANOVA with a Tukey post-hoc test. Differences were considered significant if the p-value was lower than 0.05.

## Results

### Biofilm acidogenicity

The pH decreased significantly more than the other conditions during the time of the assays when *S. mutans* was the initial enamel colonizer (Sm-Ss) and in the monospecies control Sm-Sm (p<0.05). Both conditions showed the most acidogenic potential (pH 4.5) when compared with the other groups, starting around 88 h and lasting until the end of the experimental phase (p<0.05) (Fig. 1A). Monospecies biofilms of *S. sanguinis* (Ss-Ss) and the dual-species Ss-Sm showed lower acidogenic potential, with a pH around 6.0, compared to any other condition (p<0.05). Of interest, Ss-Ss showed a significant higher pH value than Ss-Sm only after 112 h of incubation (p<0.05), making prolonged net demineralization unlikely. Interestingly, acidogenicity seems to be intermediate (pH 5.0 to 5.5) when both species are inoculated at the same time (Ss=Sm) (Fig. 1A).

**Fig. 1:**
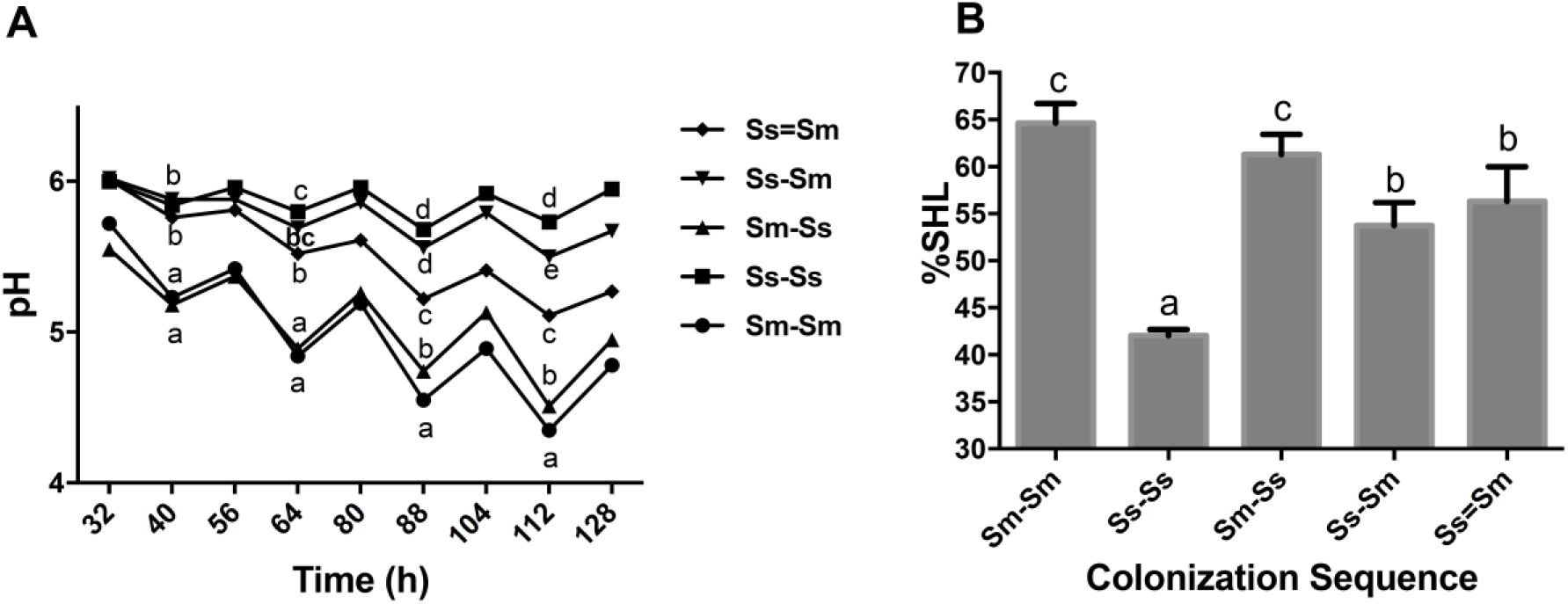
**Biofilm acidogenicity** (A). Biofilms were exposed to 10% sucrose for 5 min, 3x/day under different colonization sequences (as indicated) on enamel slabs: Ss=Sm, Ss-Sm, Sm-Ss, Ss-Ss and Sm-Sm. Medium pH was measured twice per day during the 5 days of experiment. Each point in the plot depicts mean of two independent experiments, each in triplicate (n=6). Different letters represent significant differences (p<0.05). **Enamel demineralization** (B). Enamel slabs from each biofilm exposed to cariogenic challenges with 10% sucrose were retrieved from the orthodontic wire and cleaned of the adhered biomass. Initial and final surface microhardness (SH) was measured before and after the experiment, respectively to assess percentage of SH loss (%SHL). Bars denote mean values of two independent experiments in triplicate (n=6). Error bars show the standard deviation. Different letters represent significant differences (p<0.05).

### Enamel demineralization

The percentage of surface Knoop microhardness loss (%SHL) is also influenced by the colonization sequences (Fig. 1B). Thus, the highest %SHL was observed in the Sm-Ss sequence (p<0.05) just above 60%, without differences with the Sm-Sm control biofilm, but higher than any other condition (Fig. 1B). When *S. sanguinis* was the primary colonizer (Ss-Sm), there was a significant reduction in demineralization, when compared to the *S. mutans*-primarily colonized biofilms (p<0.05), without statistical differences with Ss=Sm. However, *S. sanguinis* monospecies control biofilm showed the lowest %SHL.

Regarding the characteristics of the biofilms, there were significant variations in the properties of the different biofilms, including biomass, total protein content and insoluble extracellular polysaccharide formation (Table 1).

**Table 1:** Biofilm properties in different colonization sequence.

| Colonization sequence | Biomass (mg) | Total protein ($\mu\text{g}/\text{mg}$ biomass) | IEPS ($\%/ \text{mg}$ biomass) |
| --- | --- | --- | --- |
| **Sm-Sm** | 2.12 (0.26)<sup>d</sup> | 9.47 (1.56)<sup>b</sup> | 9.60 (3.52)<sup>a</sup> |
| **Ss-Ss** | 0.38 (0.21)<sup>a</sup> | 5.25 (0.52)<sup>a</sup> | 2.42 (1.36)<sup>b</sup> |
| **Sm-Ss** | 1.62 (0.26)<sup>c</sup> | 7.85 (0.53)<sup>b</sup> | 7.01 (1.40)<sup>ab</sup> |
| **Ss-Sm** | 0.71 (0.19)<sup>ab</sup> | 5.90 (0.66)<sup>a</sup> | 3.06 (0.99)<sup>b</sup> |
| **Ss=Sm** | 1.00 (0.35)<sup>b</sup> | 7.77 (0.66)<sup>b</sup> | 5.00 (1.97)<sup>b</sup> |

Mean (SD), n=6; IEPS: insoluble extracellular polysaccharides.
Comparisons were made vertically, for each dependent variable and among the different inoculation sequences. Different letters represent statistically significant differences ( $p < 0.05$ ).

### Biomass

When *S. sanguinis* adhered first to enamel (Ss-Sm), biofilms resulted in lower biomass than those where *S. mutans* adhered first (Sm-Ss) (p<0.05). Both single-species controls resulted in the highest (Sm-Sm) and the lowest (Ss-Ss) biofilm formation (p<0.05), respectively. Biofilms formed with *S. sanguinis* as the initial colonizer (Ss-Sm) showed lower biomass than those inoculated with both bacteria at the same time (Ss=Sm), but the difference was not statistically significant (p>0.05).

### Protein content of the biofilm

The lowest protein content in the biofilms was detected in the Ss-Sm condition and the Ss-Ss control (p>0.05). No differences were detected when *S. mutans* was the first colonizer, in the monospecies Sm-Sm control or when both species colonized at the same time (Ss=Sm) (p>0.05).

### Insoluble extracellular polysaccharide (IEPS) formation

*S. mutans* biofilms showed higher IEPS formation when compared to *S. sanguinis* (p<0.05). The lowest polysaccharide formation was detected when *S. sanguinis* was inoculated before *S. mutans*, but still slightly higher than the Ss-Ss biofilm (p>0.05).

### Bacterial counts

Viable cells counts (Fig. 2A) showed that *S. sanguinis* cells were drastically reduced when *S. mutans* was the initial enamel colonizer, compared to any other bacterial combination (p<0.05). Compared with the Sm-Sm monospecies control, *S. mutans* cells were significantly reduced in any combination when *S. sanguinis* was present as the first colonizer and even further, in the Ss-Sm and Ss=Sm biofilms (p<0.05), without differences between them (p>0.05).

**Fig. 2:**
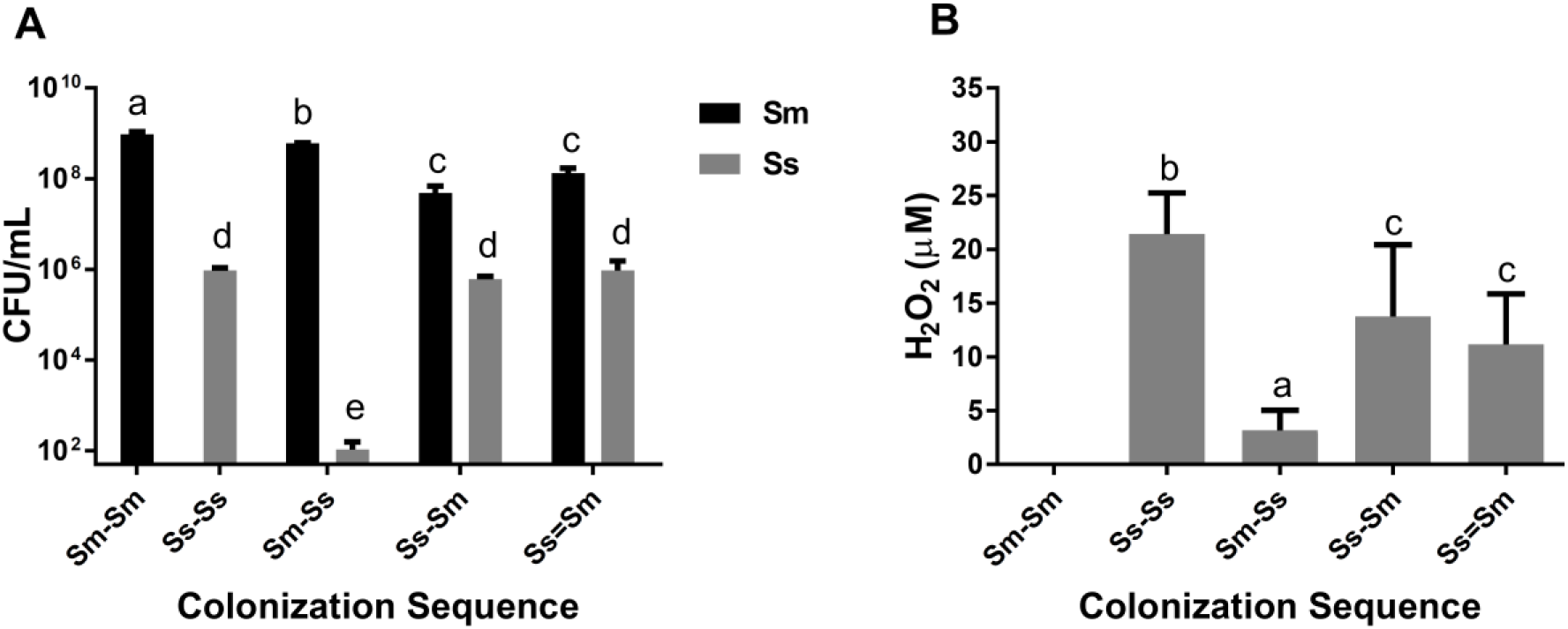
**Viable microorganisms** (A). Mean counts of *S. mutans* (black bar) and *S. sanguinis* (grey bar) expressed as CFU/mL were determined in each colonization sequence. Bars represent mean values of two independent experiments in triplicate (n=6). Error bars show the standard deviation. Different letters represent significant differences (p<0.05). **H_2_O_2_ concentration** (B). Production of H_2_O_2_ (μM) in each biofilm condition as described in Methods. Bars show mean values of two independent experiments in triplicate (n=6). Error bars show the standard deviation. Different letters represent significant differences (p<0.05).

### Hydrogen peroxide production

Despite a decrease when compared to the monospecies condition, when *S. sanguinis* was the first colonizer (Ss-Sm) or when both species colonized at the same time (Ss=Sm), there was a sustained H_2_O_2_ production (Fig. 2B). Conversely, when *S. mutans* adhered to the enamel first, a drastic reduction in H_2_O_2_ was observed (p<0.05).

## Discussion

In the present study, we modeled the dental biofilm, by confronting a commensal species; *S. sanguinis* with a cariogenic species; *S. mutans*, in a scenario where they compete for the same ecological niche. The opportunity of enamel colonization was used as the trigger for the competitive relationship, under a steady cariogenic challenge induced by sucrose. The rationale behind these studies is that, within a cariogenic environment simulated by frequent sucrose exposure, if one of the competing species colonizes first, they can mount a response to create hostile environmental conditions for the further late colonizer microorganism. Thus, *S. mutans*, in this example, can initiate and mature a cariogenic biofilm with acidic characteristics, which can exclude competitors (23).

When analyzing the acidogenicity at different times for each biofilm condition, a strong decrease in pH values (<4.5) was observed in the biofilms of *S. mutans* as single species, in addition to their highest viable counts. Conversely, Ss-Ss and Ss-Sm sequences exhibited the highest pH values (close to 6.0), which was consistent with the highest viable cell counts of *S. sanguinis*. It has been described that *S. sanguinis* is endowed with alternative mechanisms to adapt its environment and outcompete cariogenic competitors, such as *S. mutans*. For example, the arginolitic property of *S. sanguinis* is mediated by the arginine deiminase system (ADS). The ADS is able to generate ammonia, a metabolite that raises the pH and maintains it above the critical values of demineralization for the enamel (24–26). Besides, the ADS can be activated in slightly acidic conditions. This is consistent with the clinical data of *S. sanguinis* being more abundantly isolated from caries-free children (27) and adults (28).

It should be noted that the intermediate pH that was observed in the biofilms inoculated with both bacteria at the same time could indicate only moderate competition under these conditions. This is consistent with the inhibition data obtained on agar plates between both species (9). This approach with dual-species biofilms adhering on enamel and under cariogenic environments had not been previously assayed.

Despite the lack of statistical differences, when both species adhered to enamel at the same time, demineralization increased, but not to the level of the condition with *S. mutans* as the pioneer colonizer. This suggests that competition is more intense when a commensal species primarily establishes biofilms and a cariogenic microorganism attempts to colonize the niche. This is consistent with previous *in vitro* studies, showing that the inoculation sequences determine the characteristics of the oral biofilm (9).

*S. sanguinis* viable counts showed no significant differences when the enamel was first or at the same time colonized with *S. mutans* relative to its single-species biofilms. This is probably because *S. sanguinis* activates its ADS system, raising the pH and thus preventing it from being displaced from the biofilms. Coincidently, this occurs along with the lowest values of demineralization observed in the corresponding enamel slabs. Cariogenic biofilms established early on enamel by *S. mutans* have strong adherent properties, mainly due to the production of soluble and insoluble extracellular polysaccharides (23). This property makes it difficult for other less adherent cells to colonize and displace the cariogenic species. Notably, *S. sanguinis* synthesizes water-insoluble glucans, but in low amount (29).

Regarding biofilms properties, the lowest polysaccharide formation was detected in biofilms when *S. sanguinis* was the first adhering species, suggesting that *S. sanguinis* inhibits *S. mutans* colonization.

Likewise, protein and polysaccharide production followed the same trend as above, suggesting that early biofilms with *S. sanguinis* interfere with *S. mutans* colonization and the formation of thicker biofilms. When both species coexist in the dual-species biofilm, there seems to be an equilibrium in which neither manages to outcompete the other. This protective behavior may be the result of an activation of virulence factors (30, 31). The expression of virulence genes associated with these species and their molecular mechanisms have been studied. Previously, our research group analyzed the transcriptional expression of the *gtfs* genes of both bacteria and the *spxB* gene of *S. sanguinis* using the same experimental approach and design than that of this article (32). Interestingly, all genes were overexpressed when either species acted as the invading microorganism over an already formed biofilm by the antagonistic species, arguably in an attempt to colonize. Taken together, these data seem to suggest that a cariogenic environment posed by sucrose is not enough, by itself, to modify the dynamics of colonization on enamel. Although Gtf expression seems insufficient to outcompete the early colonizer, other virulent factors may be activated for competition. For example, the expression of mutacin I by *S. mutans* may act as a potent virulent factor to maintain primary colonization and avoid competition (9, 11).

The antagonism observed may also be determined by sucrose availability and the resulting acid production. As already mentioned, acidic conditions created by *S. mutans* create an hostile environment for *S. sanguinis*, inhibiting the expression of the pyruvate oxidase enzyme, responsible for the production of H_2_O_2_ (10, 33). Yet, *S. sanguinis* ADS is acid-tolerant and could contribute to maintaining H_2_O_2_ production by SpxB (34).

In this study, although *S. sanguinis* produced a smaller amount of hydrogen peroxide in the presence of *S. mutans* (and similar viable counts in Ss-Sm and Ss=Sm biofilms), there was a sustained H_2_O_2_ production creating a more competitive environment. This could explain the similar viable counts of *S. mutans* under the Ss-Sm and Ss=Sm biofilms conditions. As expected, the single species control with *S. mutans*, failed to show peroxide production, as *S. mutans* cannot produce H_2_O_2_ (14). Consistent with our results, H_2_O_2_ production by *S. sanguinis* is capable of inhibiting *S. mutans* (9, 35).

Production of H_2_O_2_ is ubiquitous among the oral commensal streptococci. *S*. *sanguinis*, however, is resistant to its own H_2_O_2_ (10, 36), which could be a key component in the maintenance of oral ecology associated with healthy conditions (34).

The results from these studies contribute to shed light on understanding the complex biological interactions in the dental biofilm under cariogenic conditions, especially when commensals are the predominant species in conditions compatible with oral health. *S. sanguinis* has been proposed as a model microorganism of molecular commensalism (13). In this context, it has been described that the expression of *spxB* is not affected by the presence of sugars (37) and the production of H_2_O_2_ is not altered by moderate pH changes (38). Thus, apparently under conditions of excess of sugars, acidic pH and *S. mutans* as a first colonizer, S*. sanguinis* cannot compete and displace *S. mutans*. Under a sucrose-induced cariogenic ecological environment, initial enamel adherence by commensal biofilms seems to induce more intense competition against a canonical cariogenic species, reducing cariogenicity (acidogenicity and demineralization). Biofilm formation with cariogenic species appears to preclude the establishment of a commensal-rich biofilms. These results must be interpreted as proof-of-principle to test novel hypothesis in a clinical setting.

In conclusion, continuous exposure to sugars seems insufficient by itself for establishing a cariogenic biofilm. Early colonization of the enamel by highly cariogenic species, like *S. mutans*, appears to be also needed. Promoting early colonization by commensal species, such as *S. sanguinis*, could help maintaining symbiosis and delaying dysbiosis.

## Author Contributions

RAG and ND conceived the idea and designed the experiments. ND performed all the experiments. ND and CL processed and analyzed the data and drafted the first manuscript. RAG and CL wrote the final manuscript. JK critically revised and contributed with new ideas to the paper. All authors revised and approved the final version of the article.

